# Automated container-less cell processing method for single-cell proteomics

**DOI:** 10.1101/2022.07.26.501646

**Authors:** Cory Matsumoto, Xinhao Shao, Marko Bogosavljevic, Liang Chen, Yu Gao

## Abstract

Single-cell genomics and transcriptomics studies enabled us to characterize cell heterogeneity in various tissues, which helped us to better understand the biological system and disease progression. Single-cell proteomics, which directly measures the protein expression level, has the potential to further enhance our knowledge by providing not only a more direct measurement but also crucial information cannot be captured by genomics or transcriptomics study, such as protein activation states and post-translation modification events. One of the main challenges of single-cell proteomics is the large sample loss during sample preparation, which is largely unavoidable using standard proteomics protocols. Protein and peptide loss to the container surface is a well-known phenomenon but often overlooked in larger-scale (>1 µg) proteomics experiments. When it comes to single-cell proteomics with only picograms of protein samples, this loss becomes non-negligible and often dictates the outcomes of the experiment. More importantly, sample processing through multiple pipette tips and containers often introduces random errors, which undermine the ability to detect true heterogenous cellular events. To solve these problems and further improve the throughput and reproducibility of the single-cell proteomics experiments, we developed an automated container-less cell processing platform, utilizing acoustic levitation to process cell samples in the air. Our platform automatically performs cell lysis, protein reduction, alkylation, digestion, and peptide labeling in the air, without any sample transfer step or container. The digested and labeled peptides are then directly injected into the capillary LC-MS/MS system for analysis, eliminating manual steps and conserving most of the sample materials for proteomics analysis. Our initial test shows at least 30% improvement in peptide signals over conventional methods. This process can be performed in parallel to further improve sample processing throughput.

## Introduction

Single-cell genomics and transcriptomics studies have enabled us to truly characterize the cell heterogeneity in various tissues and model organisms, which helped us to better understand the biological system and disease progression with unprecedented resolution.^1,2^ Being able to observe and identify cellular events at single cell level, is a major step forward for both biology and medicine. Single-cell proteomics, which relies on measuring the sample directly without any sample amplification, currently lags behind genomics and transcriptomics techniques in terms of throughput and depth.^3^ However, recent advances in the field, employing many clever methods and advanced sample preparation techniques, greatly improved the capability of single-cell proteomics, thus narrowing the gap to other omics techniques. ^4–10^ The unparalleled advantages of single-cell proteomics lay within its ability to measure protein directly, making the identification of subtle differences in protein activation states and post-translational events possible. With better sensitivity and higher throughput, it is reasonable to expect single-cell proteomics will help us to better capture rare but important cellular events in the foreseeable future.

One of the main challenges of single-cell proteomics, is the large sample loss during sample preparation, which is largely unavoidable with most of the current methods. Protein and peptide absorption on solid surfaces, e.g., glass and polymers, are well-known phenomena that have been studied extensively. ^11–14^ This is still often overlooked in larger-scale (>1 µg) proteomics experiments due to its uniform and negligible impact on protein identifications and quantitation. When it comes to single-cell proteomics with only picograms of protein samples dissolved in microliters of solvent, this absorption loss becomes no longer negligible and often dictate the outcome of the experiment. To mitigate this problem, various methods have been developed to minimize sample loss, including nanodroplet methods and better surface treatments/coatings. ^6,7^ However, without truly eliminating the liquid-solid interface, it is hard to eliminate sample loss during liquid transfer and intermediate storage steps. More importantly, sample processing through multiple pipette tips and containers often introduces random errors, which undermine the ability to detect true heterogenous cellular events.

To solve these problems and further improve the throughput and reproducibility of the single-cell proteomics experiments, we developed an automated container-less cell processing platform, utilizing acoustic levitation to process cell samples in the air (**Fig. 1, supp. video 1**). Acoustic levitation uses sound waves to create low pressure zones (nodes) in the air to trap object. It has been employed to study drugs and peptides without sample loss. It has been shown that purified peptides and small molecules can be ionized directly from the levitated droplet, making it an ideal system for small sample handling. We adopted a multi-emitter single-axis design, the TinyLev,^15^ which provides more levitation power that allows us to levitate enough liquid to perform cell processing. Our design utilized the levitation power of TinyLev and uses an enclosed system to archive a steady environment that could levitate and heat the droplet for hours. We designed automated capillary needles to deliver different reagents to perform cell lysis, protein reduction, alkylation, digestion and even isotope labeling all within the levitated droplet. The final digested and labeled peptides are then directly injected into the capillary LC-MS/MS system for analysis, eliminating liquid transfer and storage steps to conserve most of the sample materials for mass spectrometry analysis. Our initial test shows at least 30% improvement in peptide signals over microtube methods. Our system can be built in parallel to further improve sample processing throughput, making it possible to process and sample hundreds of cells per day with improved sensitivity.

**Figure 1.**
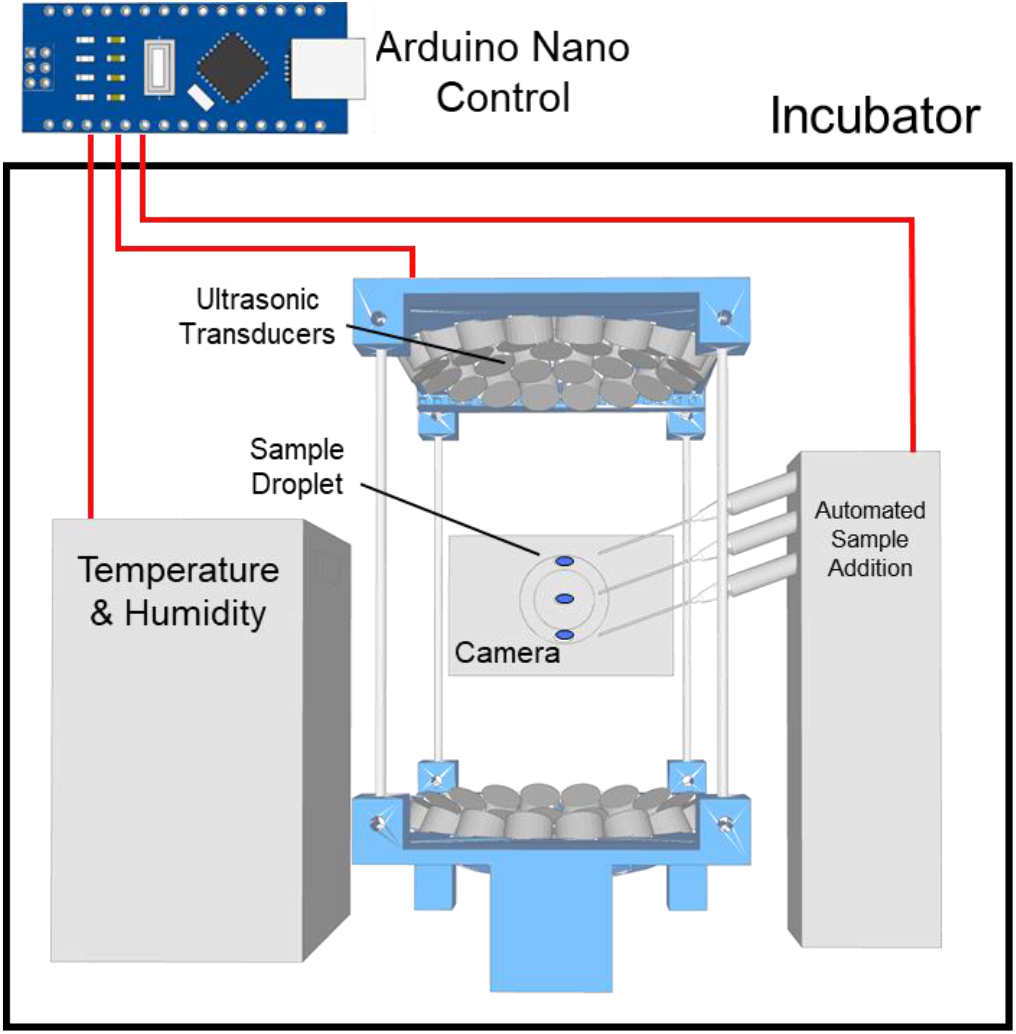
3D model of the inside of the enclosed system. An Arduino controller is used to control temperature and humidity level to keep a steady environment for the droplets. Multi-emitter single-axis acoustic levitator design adopted from TinyLev with modifications. An Arduino controlled stepper motor and syringe pumps are used to automatically deliver samples and reagents. A camera is mounted for visualize and control of the droplet.

## Results

### General design of the system

As shown in **Fig.1**, the system is designed within an enclosed system to keep the temperature and humidity stable. Our design ensures minimum airflow turbulence is introduced in all steps to maximize droplet stability (**supp. video 1-2**). An Arduino controller is used to control the ultrasonic transducers as well as other components such as humidity/temperature control and automatic sample addition arms. When a single cell is deposited as a droplet, pure water is added to perform hypotonic cell lysis. After cell lysis, the droplet is moved up to the correct position to add different reagents for digestion as needed (**supp. video 1 and 3**). Digestion usually takes 30 – 90 minutes with trypsin at 25 – 50 °C. After digestion, the top capillary connected to the automatic arm is used to aspirate the digested sample into a C-18 loading column and then injected into a nanoflow LC-MS/MS system for further analysis. Depending on the sample processing method, an additional online desalting step might be performed.

### Comparison of protein and peptide loss due to microtube storage

We first tested if our platform reduces protein loss compared to the conventional microtube method. For the comparison, we tested both PCR tubes and low-binding centrifuge tubes versus the levitated sample. Purified proteins and digested peptides from whole lysate of HEK293T cells were used for comparison to reflect the complexity of single-cell samples. The result of the PCR tubes and the low-binding centrifuge tubes are virtually identical, showing the total exposed surface area might be the most important factor in protein and peptide absorption. As shown in **Fig. 2**., no significant sample loss was observed with 200 ng of peptides, comparing the container and container-less methods. In all other comparisons, significant sample losses were observed. As we decreased the amount of sample, larger differences were seen. In 10 ng protein sample test, the container-less method recovered more than 50% more samples than the PCR tubes, showing the elimination of protein loss due to surface absorption. Limited by our detection method (micro-BCA assay), we cannot test lower than 10 ng of samples. However, we believe this trend will continue and becomes very significant when handling picograms of proteins and peptides derived from a single cell.

**Figure 2.**
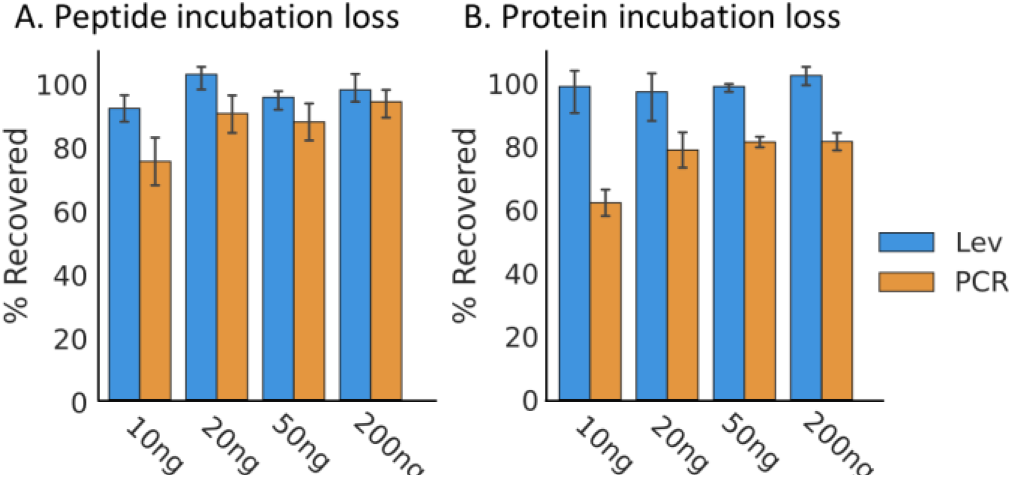
Protein and peptide incubation loss after 1 hour of incubation in PCR tube (blue) or levitation system (orange). A significant loss is observed when the amount of sample decreases to 10 ng.

### Comparison of cell processed within container and container-less methods

Single-cell proteomics probes the cell heterogeneity, and each individual cell contains very different amount of protein. A direct comparison of single cells processed by both contained and container-less methods resulted in indistinguishable effects from cell heterogeneity and the different methods. To make a fair comparison, we used 50-cell samples to compare the result of the whole proteomics sample processing pipeline. In short, 50 HEK293T cells are incubated either in a microtube or in the air using levitation system. Same amount of reagents are added to perform a standard cell lysis and digestion. After digestion, both samples are labeled the same way using TMT reagents following the SCOPE2 method. ^4,5^ In total, we identified around 700 proteins and 3,000 peptides.

We first compared the intensity distribution from the two container and two container-less samples (**Fig. 3**). The result is obvious. The two container-less sample resulted in much higher intensity throughout the entire experiment, at both protein and peptide level.

**Figure 3.**
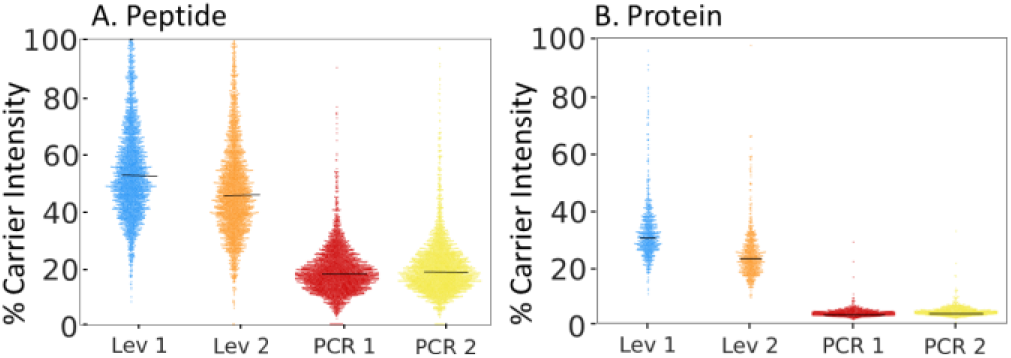
Protein and peptide intensity distribution, compared to the 200-cell carrier channel. The two container-less sample resulted in significantly higher intensity than the two container (PCR) samples.

To measure the noise level, we have also included empty channels as control. These empty channels also resulted in some signal intensity, which was used as the standard for noise. We used the average intensity of the intentionally left empty channels as the standard intensity to compare signals from the two container channels and two container-less channels. When comparing the quantitation signals of samples to empty channels, as shown in **Fig. 4**, PCR samples suffered from a significant identification loss. When we use 5x or 10x standard intensity as the filter (S/N = 5 or 10), the protein identification numbers are around 300 and 500 for the two container samples. In comparison, the two container-less sample resulted in near 700 proteins in both cases. When comparing the peptide identification, this phenomenon is even more obvious. When filtered by S/N = 10, the two container samples resulted in only 1/3 the identification comparing to the container-less samples.

**Figure 4.**
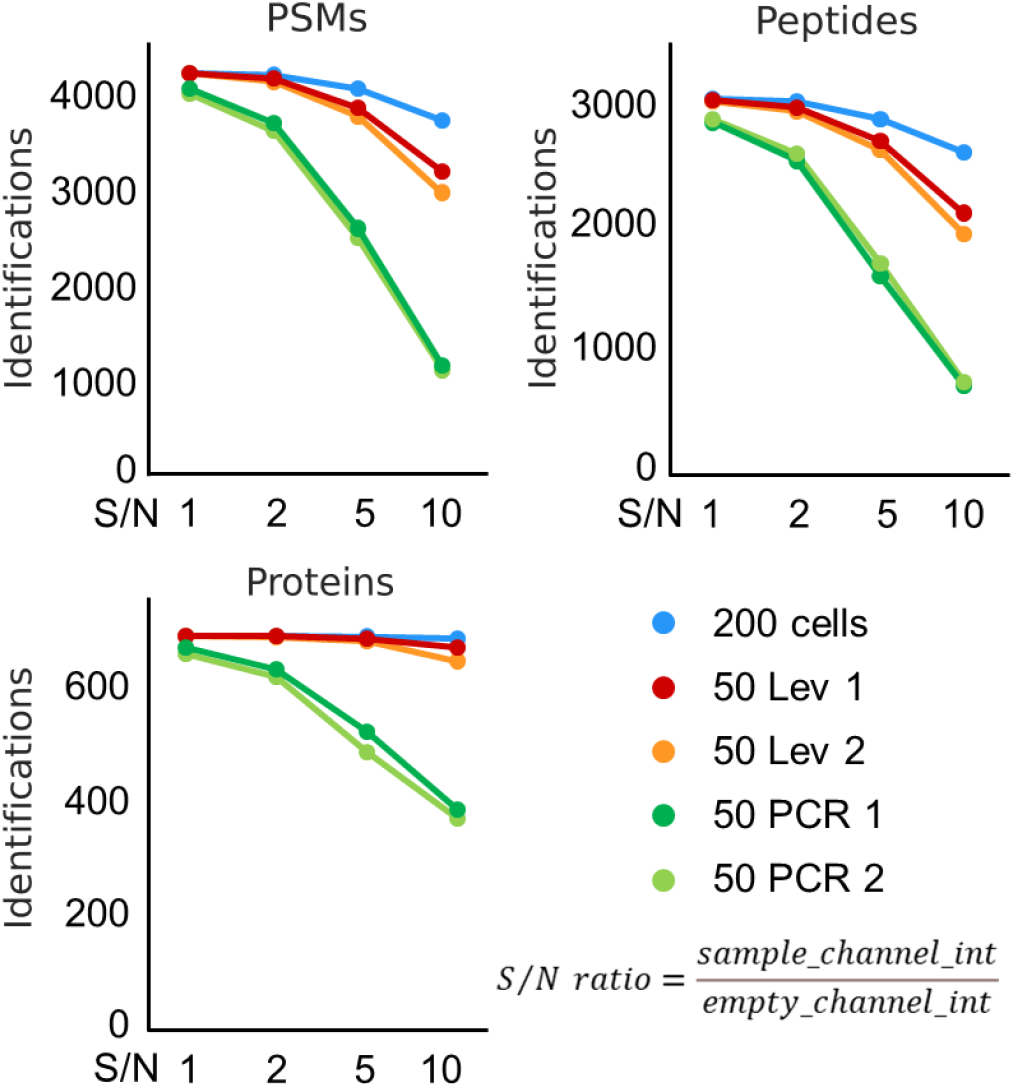
Protein, peptide, and PSM identifications at different intensity signal/noise (S/N) level. Average intensity from empty TMT channels is used as standard noise level. S/N ratio is calculated by dividing the sample intensity by the standard noise level. Container samples suffered from significant ID lost when using higher (5 and 10) S/N ratio for filtering.

### Differences of the identified peptides from container and container-less method

When we compared the average peptide length and the grand average of hydropathicity index (GRAVY) index score ^16^ of the peptides identified from container and container-less samples, we found a significant difference of distribution (**Fig. 5**). We took the log2 intensity ratio of the same peptides detected in both methods and tested for possible correlation between the peptide intensity ratio and peptide length and GRAVY scores. We used Mann-Kendall test to detect for possible trends.

**Figure 5.**
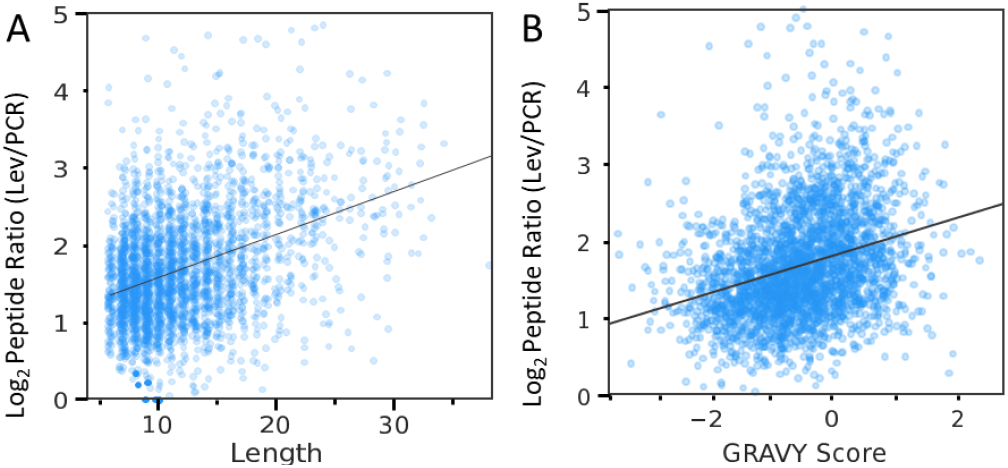
Correlation of the log2 peptide intensity ratio of container-less (Lev) / container (PCR) samples to Length and GRAVY score. There are significant trends (Mann-Kendall test, p<0.01) detected between 1. Log2 peptide intensity ratio and peptide length, 2. Log2 peptide intensity peptide ratio and peptide GRAVY score.

Both peptide length and GRAVY score exhibited significant trends (p < 0.01) with the log2 peptide intensity ratio. At the same time, the GRAVY score does not correlate with the peptide, showing the possibility of two independent factors affecting peptide absorption. This comparison shows that the plastic container selectively absorbs peptides with certain properties and length, which may result in systematic biases in single-cell proteomics analysis.

## Discussion

Here, we describe the first container-less cell processing system with acoustic levitation. We have showed by eliminating the liquid-solid interface and sample transfer steps, more protein samples could be conserved, which resulted in more detectable protein and peptide signals from the mass spectrometry. Our system is able to stabilize and process single cell as well as many cells within a levitate droplet, without the help of any container. By precisely delivering reagents with syringe pumps and capillaries, our platform is able to perform cell lysis, protein reduction, alkylation, digestion and isotope labeling of the peptides automatically. This automatic process can be parallelized by running multiple system at the same time. Our platform provides a reproducible and high-throughput way to process single-cell proteomics samples, paving the way for large-scale single-cell proteomics analysis.

## Materials and Methods

### Construction and basic operation of the Acoustic Levitator

A transducer frame and set-up were constructed using 3D printing of a modified previously reported design ^15^ controlled by an Arduino nano and powered by a DC power supply. A container was built using a modified incubator with walls lined with 0.25-inch sound and heat insulator material to optimize sound reflection and heat retention. An access point was built using a 3D printed design with magnets for ease of accessibility. A humidifier was custom built using an ultrasonic transducer and fan for the production of humidity and an induction coil controlled by a temperature probe set to a specific temperature. The automated arms of the design were controlled by an Arduino nano and assembled utilizing linear actuators with a metal rod and 3D printed frame. The aspiration and elution capillary were fashioned using a 100um I.D. deactivated silica, using an open flame to bend the glass to the desired angle.

The custom-built humidifier was filled with D.I. H_2_O, and the desired temperature was set inside the incubation container. The levitation device was placed in the container, and the voltage was initially set to 12V. After an hour of stabilization time for the temperature and humidity to build up, the voltage was brought down to 10V. The optimized location of the nodes was found by adjusting the phase of either the top or bottom array of transducers to achieve the lowest current draw of the device. From here, samples can either be added or aspirated through the direct addition of a micropipette or the automated arm.

### Protein and peptide loss assessment

To quantitatively assess the loss of proteins and peptides to PCR tubes, proteins derived from HEK lysate and peptides derived from a HEK protein digest were incubated in the levitation device or in a PCR tube. Based on the limit of detection of the quantitative assays used, quantities of peptides and proteins were assessed, totaling at least 100ng; 10 increments of 10 ng, 5 increments of 20 ng, 2 increments of 50 ng, and 1 increment of 200 ng. HEK protein was either incubated for 1hr at 45C in the levitation device or a PCR tube at a volume of 10uL before being collected in a low-binding microtube. Standards were directly transferred to a low-binding microtube. Standards and incubated samples were then dried down completely before being reconstituted by 5 µL of H_2_O with 5uL of micro-BCA assay solution and incubated at 37°C at 1hr. Proteins solutions were then quantified using a Nanodrop 100 set to 562nm based on the standard curve derived from the dried-down protein standard. HEK peptide digest was incubated in a similar way as the protein samples. Dried down peptide standards and samples were reconstituted in 10 µL of water with 70 µL of fluorescence assay solution in a black 96-well plate. Peptide solutions were quantified using a plate reader based on the standard curve derived from dried-down peptide standards.

### Comparison of container and container-less methods using 50 cells

The levitation device was set up as previously described with the temperature of the humidifier set to 65°C to ensure a constant droplet temperature of 45°C as previously recorded through direct temperature measurement through a thermocouple probe.

HEK-293T cells were harvested and diluted in 1xPBS at a concentration of 50 cells/ µL. A 200 cell carrier channel was produced using a previously PCR plate sorted by moflo Astrios cell sorter sample, which was frozen at -80C until use. The 50-cell solution was diluted with MS grade H_2_O to achieve a concentration of 50 cells/4 µL in hypotonic 0.25x PBS solution and 4 µL was either manually suspended in a levitation node by micropipette or added to a PCR tube. For the carrier channel, MS grade H_2_O was added to the 200 cells to achieve a total volume of 4 µL, and the sample was directly processed in the tube. All samples were incubated at 45°C for 15 minutes to achieve hypotonic lysis. To account for volume loss in the levitation 1.5 µL and 1 µL of trypsin solution were added to the levitation device and tube samples, respectively, to achieve a 25 ng Trypsin gold total in a 25 mM HEPES buffer solution pH 8.5. All samples were then incubated for 1 hour for digestion. Aliquots of TMT solution were dried down completely and reconstituted in MS grade H2O and immediately added to the samples at a 1 µL 11mM followed by a 45min incubation time. Unreacted TMT reagent was quenched by adding 1uL of 0.5% HA solution followed by a 30 min incubation time. The Trypsin reaction was then stopped by adding 1 µL of 5% Formic Acid to achieve a pH of ∼2. Samples were then sequentially aspirated into an in-house made c18 sample column attached to a diaphragm pump

## Author Contributions and Notes

C.M. and Y.G. designed research, C.M., X.S., M.B. and Y.G. performed research, C.M., L.M. and Y.G. performed mass spectrometry analysis, C.M. and Y.G. analyzed data; and C.M. and Y.G. wrote the paper. The authors declare no conflict of interest.

## Acknowledgments

This work is supported by NIH grant 5R35GM133416 to Y.G.

